# Jumping out of trouble: Evidence for a cognitive map in guppies (*Poecilia reticulata*)

**DOI:** 10.1101/2022.03.30.486400

**Authors:** Hannah de Waele, Catarina Vila Pouca, Dimphy van Boerdonk, Ewoud Luiten, Lisanne M. Leenheer, David Mitchell, Regina Vega-Trejo, Alexander Kotrschal

## Abstract

Spatial cognitive abilities allow individuals to remember the location of food patches, predator hide-outs, or shelters. Animals typically incorporate learnt spatial information or use external environmental cues to navigate their surroundings. A spectacular example of how some fishes move is through aerial jumping. For instance, fish that are trapped within isolated pools, cut off from the main body of water during dry periods, may jump over obstacles and direct their jumps to return to safe locations. However, what information such re-orientation behaviour during jumping is based on remains enigmatic. Here we combine a lab and field experiment to test if guppies (*Poecilia reticulata*) incorporate learnt spatial information and external environmental cues (visual and auditory) to determine where to jump. In a spatial memory assay we found that guppies were more likely to jump towards deeper areas, hence incorporating past spatial information to jump to safety. In a matched vs. mismatched spatial cue experiment in the field, we found that animals only showed directed jumping when visual and auditory cues matched. We show that in unfamiliar entrapments guppies direct their jumps by combining visual and auditory cues, while in familiar entrapments they use a cognitive map. We hence conclude that jumping behaviour is a goal-directed behaviour, guided by different sources of information and involving important spatial cognitive skills.

## Introduction

Mobile animals need to navigate to locate resources, find mates, or return to their home area, while simultaneously minimizing predation risk (Braithwaite & Girvan, 2003; Noda, Gushima, & Kakuda, 1994). Animals rely on a range of spatial cognitive skills to perform these tasks; they can recognize complex cues, form long-term memories, and regain their bearings after disorientation. For example, Clark’s nutcrackers (*Nucifraga columbiana*) can memorize the locations of thousands of caches up to nine months after caching (Balda & Kamil, 1992), and displaced honeybees find their hive by using spatial features learned during a single flight (Degen et al., 2016). Some fish species also have excellent spatial learning and memory abilities (Teyke, 1989; Warburton, 2003). Atlantic salmon (*Salmo salar*), goldfish (*Carassius auratus*), fifteen-spined sticklebacks (*Spinachia spinachia)*, and corkwing wrasse (*Crenilabrus melops)* can locate a food reward using visual landmarks (Braithwaite, Armstrong, McAdam, & Huntingford, 1996; Hughes & Blight, 2000; López, Broglio, Rodríguez, Thinus-Blanc, & Salas, 1999; Salas et al., 1996), even when released in an unfamiliar spot (Rodriguez, Duran, Vargas, Torres, & Salas, 1994). There is also evidence that some fishes (e.g. Mexican cave fish (*Astyanax mexicanus*), goldfish (*Carassius auratus*), and butterflyfishes (family Chaetodontidae) use cognitive maps to navigate their surroundings (Burt De Perera, 2004; Reese, 1989; Rodriguez et al., 1994). Cognitive maps are internal map-like representations of relationships between different landmarks in their environment, created through spatial learning (Gould, 1986; Toates, 1980; Tolman, 1948). Such an internal representation of the environment allows animals to adopt novel routes from previously unvisited points, removing the requirement of learning specific routes (Mazmanian & Roberts, 1983; Morris, 1981; O’Keefe & Conway, 1978; Suzuki, Augerinos, & Black, 1980).

To further understand mechanisms of spatial orientation, researchers have taken advantage of aerial jumping behaviour in fishes since it presents a clear directional choice of movement (Buo et al., 2020a). Aronson (1951, 1971) and White & Brown (2014) used this behaviour to demonstrate that gobies have a cognitive map of their habitat as they can leap from one tidal pool to another even when they have no visual contact before leaving the water. Guppies (*Poecilia reticulata*) also show aerial jumping (Soares & Bierman, 2013) and may therefore possess similar spatial abilities. In Trinidadian rivers, the guppies’ native range, seasonal changes and weather events connect and disconnect pools; connecting them at high water and isolating them from the main river as the water recedes. By being able to jump out of such pools, guppies should be able to return to the main river, escaping isolation and potential death as the pools desiccate. Soares & Bierman (2013) documented guppies generally starting a jump with a slow and preparatory backwards movement, followed by a forward thrust, propelling the fish into the air. These small fish are able to jump approximately 3.5 times their body length in height, with a positive correlation between jump height and back-up distance (Soares & Bierman, 2013). This suggests that guppies can actively alter jump height and likely also distance. Additionally, the preparatory phase suggests jumping is not purely reflexive, but involves a clear motive. However, whether guppies are able to direct their jump towards a specific location, and what information they base their jumping direction on, remains untested.

As guppies are capable of using spatial information in maze-experiments (Burns & Rodd, 2008b; Chapman, Ward, & Krause, 2008; Lucon-Xiccato & Bisazza, 2017b, 2017a; Miletto Petrazzini, Bisazza, Agrillo, & Lucon-Xiccato, 2017), we tested if and how guppies base their jumping directions on spatial information in two experiments. In the laboratory, we tested whether guppies use the memory of specific landmarks (cognitive maps) to jump out of ‘trouble’ in a known small water body. We used water level changes in combination with entrapment in desiccating pools to test if guppies remember which areas are safe to jump to. In this relatively small arena, the shallow areas become dry land and the deep areas remain water-filled pools. We predicted that if guppies memorize their environment and incorporate that spatial information, they should aim their jump towards the pools.

In a second experiment in the wild, we tested whether guppies show directed jumping behaviour when entrapped in unknown small water bodies. Such a situation may occur during flash floods when a fish gets washed ashore and finds itself in a desiccating body of water that lies outside its known area. We hypothesized that, in the scenario where the fish has no previous information about the specific environment, they should use general landmarks like visual and/or auditory cues. Hence, this scenario excludes the possibility of basing jumping direction on cognitive maps, and requires them to rely on external, general landmarks. For instance, the open canopy may function as a general landmark as it can indicate the middle of the river. Sunlight and gap size in the tropical forest determine different light environments, with canopy openness decreasing from the centre of the stream to the riverbanks (Endler, 1992). The canopy structure has various visual aspects that the guppies could use for orientation, including the increased light level in the middle of the river. There is evidence that several fish species show the ability to orient themselves towards light sources (phototaxis) (Barki, Zion, Shapira, & Karplus, 2014; Goodyear & Ferguson, 1969). In addition to phototaxis, the trees at either side of the river can function as a general visual landmark on which fish orient to find the middle of the stream. Although the canopy is distant, fishes have been shown to use external, general cues outside of the water to orient themselves (Rodriguez et al., 1994). Hence, the various visual aspects of the canopy structure can function as a general visual landmark on which fish orient to find the middle of the stream. Another general landmark may be the sound of flowing water as a loud local riffle may indicate flowing water and therefore a safe jumping direction. Sound can be important for orientation in aquatic animals because of its highly directional nature (Hawkins & Popper, 2018). For example, sound is a meaningful orientation and attraction cue for larvae of reef fishes (Montgomery, Jeffs, Simpson, Meekan, & Tindle, 2006; Simpson, Meekan, Montgomery, McCauley, & Jeffs, 2005; Tolimieri, Jeffs, & Montgomery, 2000). Since the sound of a riffle indicates a ‘safe’ direction towards water, it may trigger the guppies to jump towards that direction. We investigated their jump direction in a cue-conflict/cue-matching experiment, where we exposed guppies to different spatial combinations of riffle sounds and canopy openness. This experiment tested which of two general landmarks, sound or canopy openness, guppies base their jumping decision on. We hypothesized that both cues are important and predicted that guppies will show the highest directional preference when both cues match.

## Materials and Methods

We aimed to test the hypotheses that guppies can (1) use a cognitive map based on learnt spatial information, and (2) can incorporate external environmental cues (visual and auditory) to determine where to safely jump. To do so, we ran two experiments.

### Experiment I - Testing for a cognitive map in the laboratory

In a laboratory experiment, we aimed to understand whether guppies use cognitive maps to direct their jumps. The study was carried out in the animal facilities at Wageningen University and Research in April 2020. Guppies used in the experiment were 20 adult females; laboratory-reared descendants of fish collected from a high-predation population from the Aripo river in Trinidad. Prior to experiments, fish were housed in same-sex groups of six in 4 litre flow-through tanks at Wageningen University. Temperature was kept at 24°C with a 12:12h dark/light cycle. Fish were fed twice daily on seven days a week with flake food and freshly hatched *Artemia salina* nauplii.

Two identical experimental tanks (70 × 35 × 40 cm) were filled with gravel and shaped into four quadrants of different depths (two shallow areas, two deep areas; Figure 1). Two-centimetre ridges separated the quadrants from each other. Additionally, two markers, consisting of wooden sticks, were used to mark the division between the shallow and deep areas and to serve as a visual landmark for the fish. The water level in the experimental tank could be altered by siphoning out water or adding water through two identical tubes attached to the sides of the tank. The shallow areas were approximately five cm deep at high-water. At low water level, the shallow areas fell dry. The deeper areas consisted of a slope, starting its decline at the border with the shallow areas. During high water the deep area ranged from 5 to 17 cm in depth. During low water, the deepest area was 12 cm deep. In the middle of the tank there was a circular testing arena with a diameter of 9 cm that was surrounded by a 1.5 cm high ring of gravel so that the water level of the rest of the tank was not visible from the inside the testing arena during low water levels. The circular testing arena had a depth of approximately 3 cm at low levels, imitating an isolated tidal pool. A camera was mounted above the experimental tank. Both deep areas were equipped with a filter during the habituation period of the test, and stones were used in the corners of the shallow areas to limit access to the tubes used to alter the water level.

**Figure 1.**
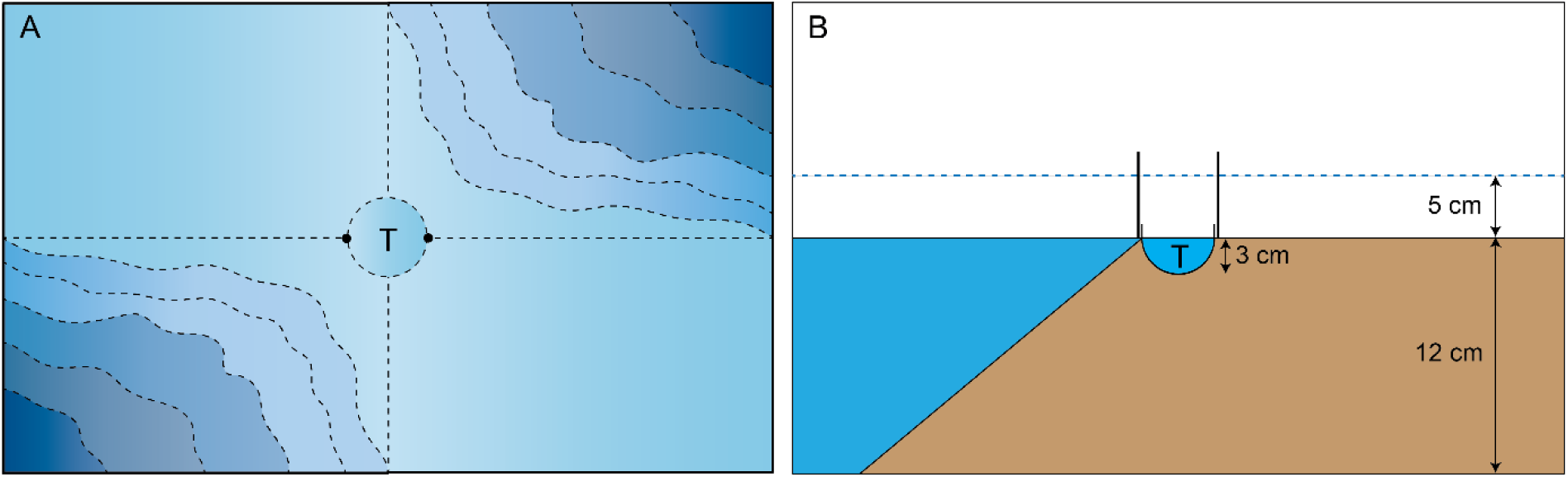
Set up of the experimental tank: view from above (A), and view from the side at low water (B) with depth representation. The experimental tank was filled with gravel to create four quadrants of different depths (two shallow areas, two deep areas). Two-cm ridges separated the quadrants from each other. Different shades of blue represent different depths, with darker tones marking the deeper areas. The “T” marks the testing arena.

Ten females were simultaneously transferred into one of the experimental tanks at high water and allowed to habituate for one hour. This and all subsequent procedures were done in parallel in the two experimental tanks. The fish were exposed to the experimental tanks for three consecutive days. They then experienced three 2-hour periodical flood cycles per day. Within one cycle, the water receded for approximately 25 min. After a 10 min rest period, the water started rising for 25 min. This was repeated twice per cycle. After finishing the third cycle the fish were fed. The same procedure was repeated on the second day. On day 3, after one full cycle, the water level was lowered once more, and the guppies were caught and moved to temporary holding tanks. Guppies were caught individually with a net, then transferred into the circular testing arena with a cup, and observed for their jumping behaviour. If a fish did not jump within 10 min, they were removed from the testing arena and returned to their holding tank and excluded for further testing. Latency to jump and landing location were determined from video footage. Note that before the testing phase the shallow areas fell completely dry at low water, yet, during the testing phase, a depth of 1 cm was maintained to avoid fish jumping on dry land and possibly injuring themselves. Thus, even when they jumped towards the shallow areas, they landed in shallow water. The water level was well below the visible threshold from within the testing arena.

### Experiment II - Testing for general spatial cues in the wild

In a field experiment, we aimed to understand whether guppies use auditory cues of moving water and/or visual cues of the open canopy to orient themselves when jumping. To do so, we conducted an experiment in the lower Aripo river in Trinidad (Lat.: 10.6667; Long.: −61.2285) in March 2020. This site was chosen due to its steady guppy population, its pronounced network of riffles and pools, and a relatively high canopy openness in the middle of the river. A riffle was defined as a small waterfall or a localized fast-flowing current generated by numerous boulders within the river (Figure S2). Apart from the sound of the riffles, the site was calm and the riffles were spaced apart, limiting the sound interference between different riffles.

We selected three locations towards the left bank and two locations towards the right bank so as to vary the direction to the open canopy. At each of these locations, two different spatial arrangements (treatments) relative to two cues were created on either side of the riffle (Figure 2). The riffle was either spatially matching – at the same side of the open canopy – or contrasting – at the opposite side of the open canopy (Figure 2). Water depth ranged from a few centimetres close to a riffle to 1 m in the deeper areas. Note that all locations were within the stream for ethical reasons. Thus, all jumping directions led back to water.

**Figure 2.**
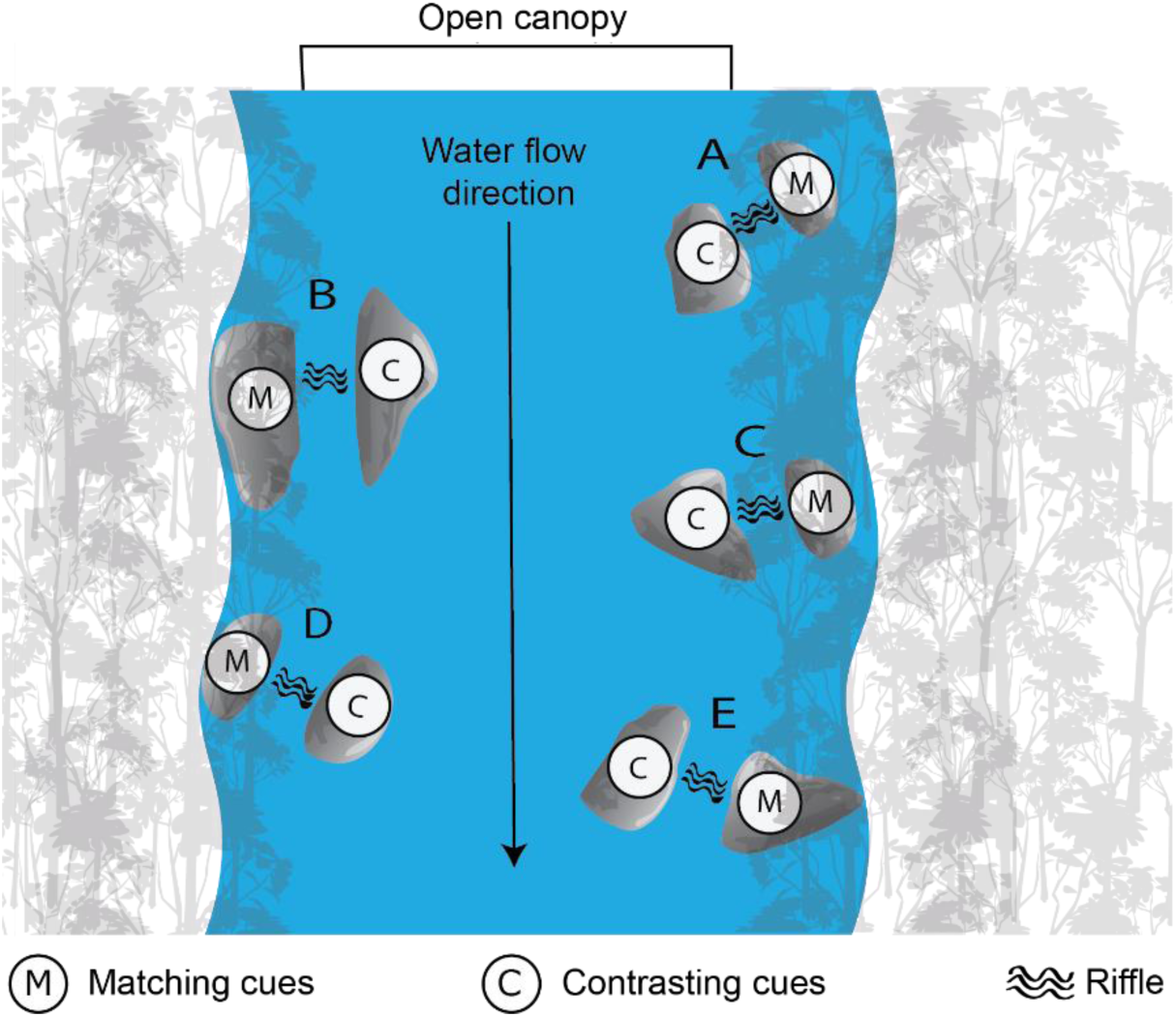
Schematic overview of the study site. Test locations are marked A through E. At each location both treatments (M: matching cues; C: contrasting cues) were tested.

Nine trials were done at two locations, and eight trials were done at the other eight locations. Female, male, and juvenile guppies (N = 492) used in this experiment were collected with dip and butterfly nets within a range of twenty meters around the test site in pools and the main stream. We chose to assay females, males, and juveniles to obtain a more representative sample of jumping behaviour in the species, but made no specific predictions on how females, males, and juveniles should differ in their jumping behaviour.

During jumping trials, guppies were placed in opaque white Styrofoam cups (5.5 cm diameter, 3.5 cm height) that were filled with 30 ml of fresh stream water so that the height difference between the water surface and cup edge was roughly 1 cm. At the start of a trial, six cups were placed near the edge of a boulder at the riffle side and guppies were individually placed into the cups. Trials were filmed for 10 minutes using an overhead stationary GoPro camera, mounted on a PVC tube structure (Figure S2). Observers kept a distance of at least one meter.

For each trial, individual fish were scored for whether or not they jumped within 10-minutes and the jump latency was recorded. Subsequently, for the fish that jumped (n = 275), still frames were isolated from the video footage and analysed to estimate the jumping angle (Fig. S1). A first frame was selected when the fish started to protrude above the water surface. In this frame, a line was drawn from the fish’s centre to the riffle centre. This line was used as a reference and was set at 0 degrees using modular arithmetic. Thus, all subsequent calculated angles were relative to this line. A second frame was then selected where the fish crossed the edge of the cup. In this frame, a line was drawn from the fish’s centre when starting the protrude to the fish’s centre when crossing the edge of the cup. The angle between the two drawn lines was calculated by measuring the shortest clockwise path using the Protractor feature in the PicPick software (version 5.0.7; 2020). This angle is referred to as the jumping angle. Three jumps were excluded from the analysis since the angle was impossible to measure due to low quality footage. The jumping angle was calculated in 272 individual guppies by a single researcher to avoid observer differences.

### Statistical analysis

#### Experiment I

To test whether the jumping rate towards the pools was significantly different than random chance we used a two-sided binomial exact test. Subsequently, to test whether the latency to jump varied between the fish jumping towards deep areas or the fish jumping towards the shallow areas we used a Wilcoxon rank sum test.

#### Experiment II

To test for differences in Jumping frequency, we used a linear mixed model (binomial distribution) with Treatment (opposite or matching cues), Life-stage (male, female, juvenile) and the interaction of Treatment x Life-stage as predictor variables, as well as a random factor for Location. We used the ‘lme4’ package (Bates, Mächler, Bolker, & Walker, 2014). Model terms were tested for significance using the ANOVA function in the car package (Fox & Weisberg, 2018), specifying Type III Wald chi-square tests. If Life-stage was a significant predictor in the model, we assessed Tukey corrected multiple comparisons between Life-stage levels using the ‘lsmeans’ function in the ‘emmeans’ package (Lenth, Singmann, Love, Buerkner, & Herve, 2019). The interaction between Treatment and Life-stage in this model was removed from the model as it was marginally non-significant and uninformative for our research question (p = 0.0506).

To test for differences in Jump latency, non-jumpers were excluded. Subsequently, we used a linear mixed model (normal distribution) with Treatment, Life-stage, and the interaction of Treatment x Life stage as predictor variables, as well as a random factor for Location using again the ‘lme4’ package. Model terms were tested for significance using the ANOVA function in the car package (Fox & Weisberg, 2018), specifying Type III Wald chi-square tests. The interaction between Life-stage and Treatment in this model was removed from the model as it was non-significant and uninformative for our research question (p = 0.403). If Life-stage was a significant predictor in the model, we assessed Tukey corrected multiple comparisons between Life-stage levels using the ‘lsmeans’ function in the ‘emmeans’ package (Lenth et al., 2019). To meet the homogeneity and normality assumptions, we performed a log (10) transformation on Jump latency.

To investigate the role of sound and/or canopy cover for jumping orientation, we used circular statistics techniques. The Rayleigh test was used to assess the uniformity of the jumping direction. If no clear direction is preferred, then all directions would be approximately equally covered on the circumference of a circle, visible in a circular uniform distribution. Any deviation from a circular uniform distribution would be indicative of a preferred direction. The Rayleigh test assumes that the data follow a unimodal and symmetrical von Mises distribution (Landler, Ruxton, & Malkemper, 2019). To rule out multimodal departures and unknown symmetry, the Hermans-Rasson test was performed. All statistical analysis were performed in R-3.6.2 (r Development Core Team 2012).

## Results

### Experiment I

In the laboratory experiment testing if guppies direct their jumps using a cognitive map, all female guppies jumped out of the testing arena. Of the twenty tested animals, 16 jumped towards a deep area and 4 jumped towards a shallow area (Figure 3A). Thus, fish were more likely to jump towards the deeper pools than expected at random with a 50:50 chance (N = 20; Z = 16; p = 0.006).

**Figure 3.**
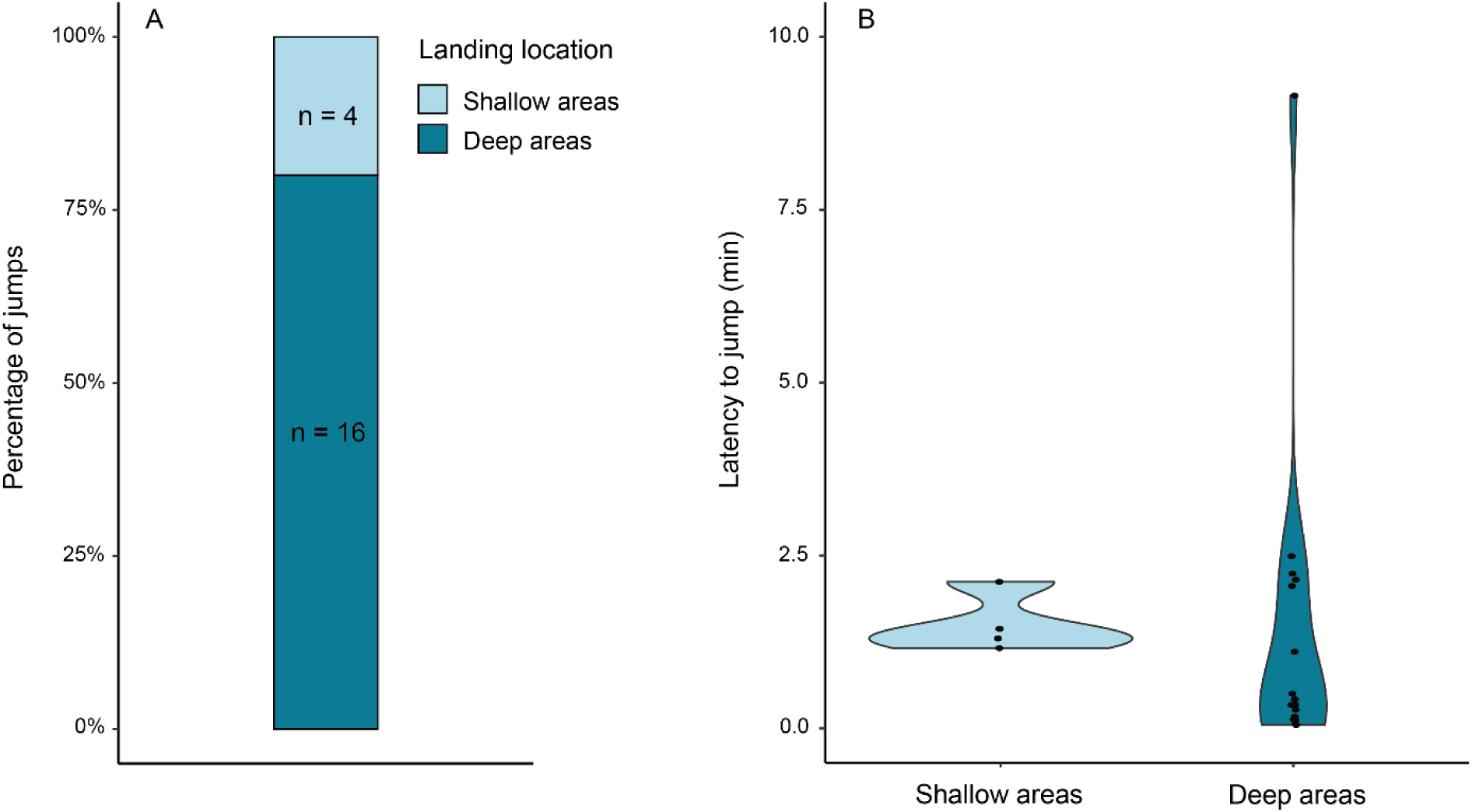
Percentage of jumps (A) and latency to jump (B) for guppies jumping into the deep areas (dark blue) and onto the shallow areas (light blue). Black markers in B show individual data points.

We found no difference in the latency to jump between the guppies jumping towards the deep areas compared to the guppies jumping towards the shallow areas (Wilcoxon rank sum test; N = 20; W = 45; p = 0.237; Figure 3B). Note that there were only 4 fish in the group jumping towards the shallow areas, thus, there is little power to assess differences in the latency to jump towards these different areas.

### Experiment II

In the field experiment testing for general spatial cues, 55.9% (275/492) individuals jumped out of the testing cup, while 44.1% (217/492) did not jump within the 10-minute experimental time. We found no differences in jumping frequency between the treatment groups (χ^2^_1,270_ = 0.019; p = 0.892), but jump frequency differed between juveniles, females and males (χ^2^_2,270_ = 23.392; p < 0.0001). Pairwise comparisons showed that juveniles jumped significantly less than males and females (Juvenile – Male: p = 0.019 and Female – Juvenile: p < 0.0001).

When filtering the data to only fish that jumped, we found no differences in the latency to jump of fish from the matching cues treatment compared to those from the contrasting cues treatment (F_1,270_ = 3.626; p = 0.057; Figure 4A). Yet, we found that juveniles, females, and males differed in jump latency (F_2,270_ = 23.229; p < 0.0001; Figure 4B). Pairwise comparisons indicated that females jumped approximately 26 seconds faster than juveniles and 19 seconds faster than males (Female – Juvenile: p < 0.0001 and Female – Male: p = 0.012).

**Figure 4.**
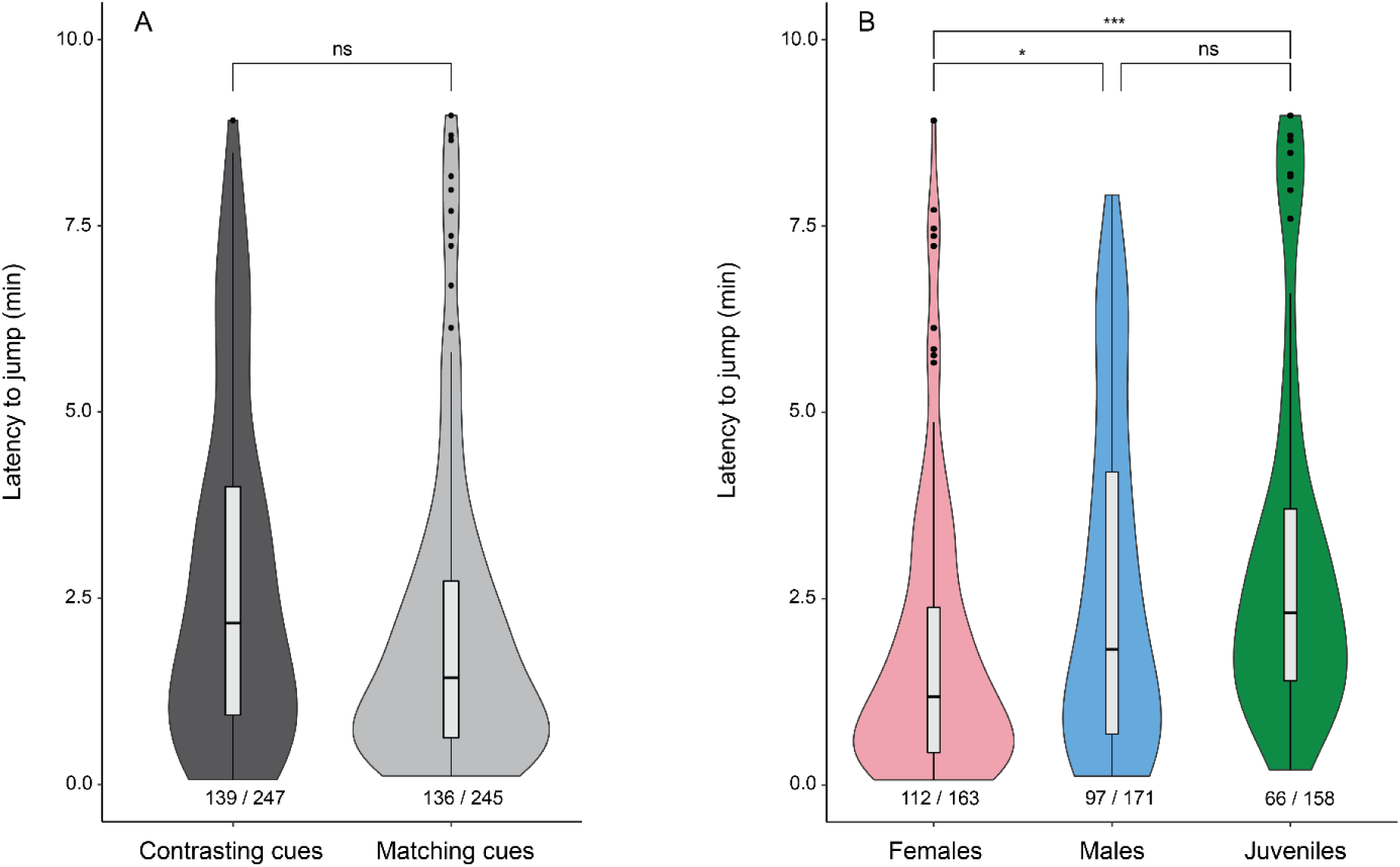
Latency to jump for the different treatments (contrasting cues and matching cues) (A) and for females, males, and juveniles (B). The proportion of jumpers for each group is noted underneath the violin plot. Horizontal lines indicate medians, boxes indicate interquartile range, and whiskers indicate all points within 1.5 times the interquartile range.

When exposed to matching cues, the jumping direction for all fish showed a significant deviation from a uniform distribution (*R* = 0.253; Z_135_ = 8.659; p = 0.0002; Figure 5A, which indicates that fish preferentially jump towards a specific direction. We found that the mean direction of the jump was 18.75 degrees relative to the middle of the riffle, pointing towards the riffle and canopy openness. In contrast, fish exposed to contrasting cues showed a randomly distributed jumping orientation, with no clear preferred jump direction (*R* = 0.058; Z_137_ = 0.463; p = 0.630; Figure 5B).

**Figure 5.**
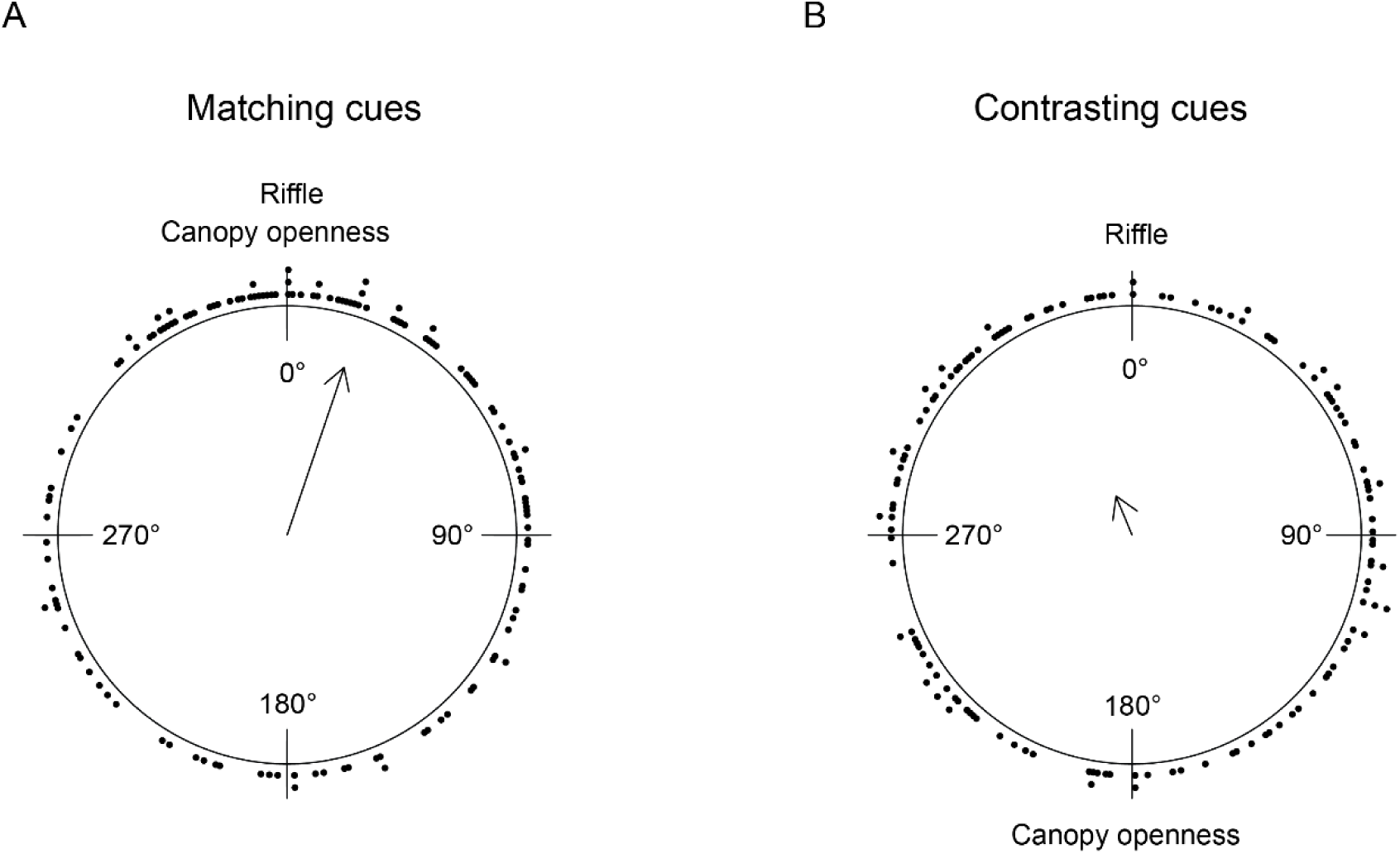
Jumping direction distribution for (A) matching cues (n = 136) and (B) contrasting cues (n = 139). The arrow points towards the mean angle and its length reflects the concentration of data. Black dots indicate individual data points.

When investigating the jump direction of males, females, and juveniles separately, only females in the matching cues treatment showed a preferred direction towards the two cues, departing from a uniform direction (Rayleigh test; 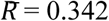; Z_60_ = 7.037; p < 0.0009; Figure 6A). Although the mean jumping direction for the males and juveniles appeared qualitatively similar (Figure 6A), the jumping direction did not differ from uniformity. When exposed to contrasting cues, neither of the sexes nor juveniles showed a preferred jumping direction (Figure 6B).

**Figure 6.**
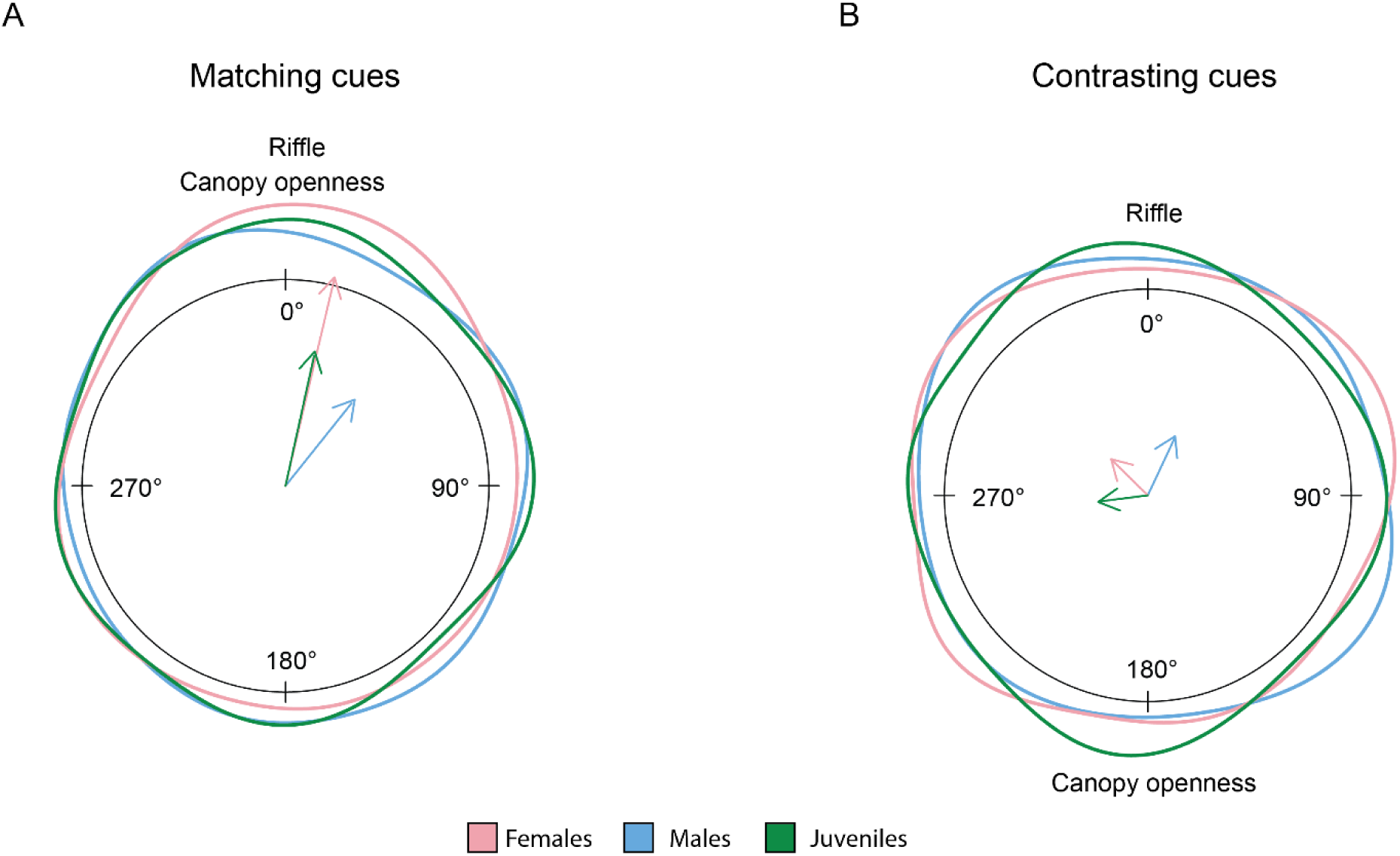
Jumping distribution for the different stages (female, male, and juvenile) for matching cues (A) and contrasting cues (B). The arrow points towards the mean angle and its length reflects the concentration of data.

## Discussion

In two complementary experiments, we found evidence that guppies direct their jumps towards a specific direction and use both a cognitive map and general landmarks to orient themselves. When guppies were confined in a small and shallow pool in a familiar environment, they swiftly jumped out to land in deeper water bodies. In contrast, when they found themselves in unknown confined water bodies, they were less likely to jump, but when they did, they oriented themselves using general landmarks such as the sound of moving water and canopy openness.

The first aim of this study was to test whether guppies have the capacity to memorize their surroundings and use this information to determine their jumping direction. We found that all animals jumped out of the test arena within minutes and most of the animals jumped towards areas with deeper water, supporting our prediction that guppies can use cognitive maps to direct their jumps. Spatial cognition has been well studied in guppies (Burns & Rodd, 2008a; Kotrschal, Corral-Lopez, Amcoff, & Kolm, 2015; Lucon-Xiccato & Bisazza, 2017c). For example, they can learn to navigate a complex maze consisting of up to six consecutive T-junctions (Lucon-Xiccato & Bisazza, 2017a). Our data adds cognitive maps to this portfolio of spatial abilities because, similar to fishes such as the amphibious spotted rock skipper and some gobies (Aronson, 1971; Buo et al., 2020b; White & Brown, 2014), they reliably jump to favourable areas even when the direct line of sight is blocked. Due to the way our test pool was designed, the guppies could not see the water level outside of the pool during the test and could only orient using the provided visual cues and/or internally stored information.

External visual cues included the quadrant division markers, cues from the experimental room, and differences in ceiling colour. While maze experiments show that guppies typically do not rely on subtle cues when navigating (Lucon-Xiccato & Bisazza, 2017a), they could have done so in this study and memorized external landmarks may be part of the cognitive map of the environment. In line with our predictions, we found that guppies have the ability to create cognitive maps of their environment and retrieve those to direct their jump in order to successfully return to deeper bodies of water.

The second experiment tested how guppies react to being trapped in small water bodies that are unknown to them but lie within their known home range. In these cases, they likely had no memory of the quality of the environment they jumped towards. We hypothesized that guppies then may rely on external visual and/or auditory cues, i.e. the canopy structure and a local riffle, to orient themselves when jumping out of an isolated area. In line with our predictions, we found that only when visual and auditory cues were aligned guppies showed clear directional jumping towards the middle of the stream, while under contrasting cues no clear directional choice was evident. Guppies seem to apply multimodal orientation as several cues need to align for effective re-orientation.

Most spatial task studies in fish have focused on the relevance of learned landmarks for orientation (e.g. Buo et al. 2020b; Odling-Smee and Braithwaite 2003). Guppies in the present study were predicted to orient spontaneously without training or reinforcement. Guppies live in large and highly changing habitats. Hence, the use of cognitive maps might not always be feasible or optimal. General landmarks, such as the geometry of the environment, should provide a better tool to orient than local landmarks when stranded in an unknown area. Indeed, the shape of the environment is crucial for reorientation in fishes, rats, and pigeons, showing that large-scale geometry of the environment is more relevant than local non-geometrical cues (Cheng, 1986; Kelly, Spetch, & Heth, 1998; Sovrano, Bisazza, & Vallortigara, 2002).

In a guppy’s habitat, we hypothesized that canopy openness and riffle sound may be suitable general landmarks. Brighter light conditions can be indicative of the location of the middle of the stream, as the canopy is less dense there. This should provide information about where the water is likely the deepest. On top of light level, the canopy also provides other visual features that animals have been shown to use for orientation, including specific shapes in the canopy pattern or the overall contrast between the trees at both sides and the open sky. For instance, ants orient using the high contrast cues created by canopy patterns (Ehmer, n.d.; Hölldobler, 1980) and in fishes, depending on species, positive or negative phototaxis are widespread (Barki et al., 2014; Forward Jr, Horch, & Waterman, 1972). Additionally to optical features, we hypothesized that water murmur, or the disturbance caused by water flowing against rocks at riffles, might function as an general auditory landmark, as fishes can use sound as an orientation cue (Montgomery et al., 2006). We found that guppies showed only a preferential jumping direction when the two hypothesized landmarks, open canopy and riffle sound, were aligned. We conclude that both general landmarks are important for orientation and are used collectively.

Several non-mutually exclusive hypotheses exist to explain aerial jumping in guppies, from predator evasion (Seghers, 1973) to means of dispersal (Soares & Bierman, 2013). Here we show that guppies can use aerial jumping to escape isolation. Streams in Trinidad, the guppy’s environment, are highly dynamic with rapid water level changes, and getting trapped in a desiccating pool after a flood or a dry period is likely a common event. While staying in these isolated pools would mean isolation or even death, uncertainty about the surroundings may prevent the guppy from jumping into an unknown environment. Costs of erroneous jumping decisions include landing on dry land with the risk of desiccation. Even though repeated flapping on dry land can help getting back into a water body, impact on the rocks may still cause physical injury. We found a clear difference on whether guppies jumped or not out of confinement between our two experiments. While all animals rapidly jumped from the known pools in the laboratory, almost 40% of animals did not jump out of unknown confined water bodies within the experimental time. We interpret this difference as evidence for a divergent cost and benefit trade off as jumping out of an isolated pool of a known environment carries less risk than jumping in a novel environment.

We also found differences between female, male, and juvenile jumping behaviour. Juveniles jumped less than adult fish, and females jumped faster than juveniles and with a stronger directional choice. While we had no clear *a priori* prediction about how groups may differ in jumping behaviour, we speculate how this difference may be explained. Divergence in risks of jumping may be one explanation; for juveniles, the risk of jumping may be greater, since a smaller body is related to higher desiccation and injury risk in fish (Odeh, 2002). Moreover, juveniles might lack landscape knowledge and jumping skills, which increases the chance of hitting a rock. Differences in experience may also explain the difference in latency time of jumping between males and males. Females in the wild are less predated on and live longer than males (Reznick and Bryant 2007; Reznick et al. 1996). Due to their increased lifespan, they can gather more landscape knowledge and jumping skills. In contrast, males, having a shorter lifespan, might have evolved to invest more in mating efforts than in jumping and orientation skills. This reasoning is supported by our findings that while all fish showed the same orientational pattern when exposed to matching cues, only females significantly favoured one direction over another. However, this line of argument seems to oppose a previous study that found that in mazes male guppies have better spatial abilities (Lucon-Xiccato & Bisazza, 2017c). The authors suggest that because males have larger home ranges and disperse further (Croft et al., 2003; Croft, Krause, & James, 2004; Darden & Croft, 2008), they have evolved better spatial cognition. Additional data is clearly needed to clarify how maze performance relates to orientational skills in the wild, focusing in particular on jumping behaviour.

To conclude, this study contributes to our understanding of how fish are able to use different cues to find suitable water bodies. In a lab experiment, we found that guppies were more likely to jump towards deeper areas when they are in a familiar entrapment, hence incorporating past spatial information to jump to safety. In a matched vs. mismatched spatial cue experiment in the field, we found that animals only showed directed jumping when general visual and auditory cues matched. Hence, in this study, we show that fish rely on the use of cognitive maps and a combination of general landmarks to orient themselves during aerial jumping.

## Acknowledgements

We thank Amy Deacon for assistance during the field trip in Trinidad, and Bart Pollux for providing stock fish to start our experimental populations at WUR. We further thank the staff from the BHE group and Carus at WUR for technical and husbandry assistance.

## Contributions

C.V.P., R.V.-T., D.J.M., D.v.B., E.L., L.M.L., and A.K. conceived the study. C.V.P., R.V.-T., D.J.M., D.v.B., E.L., and L.M.L. collected the data. H.D.W, D.v.B., and E.L. analysed the data and wrote the first version of the manuscript. All authors provided feedback on earlier versions of the manuscript and contributed to its final version.

## Ethical considerations

The care and use of experimental animals complied with animal welfare laws, guidelines and policies as approved by the Animal Welfare Body (Instantie voor Dierenwelwijn, IvD) at Wageningen University and Research and by the Fisheries Division of the Ministry of Agriculture, Land and Fisheries of Trinidad and Tobago. Experiments involved behavioural observations, and stress to the animals was minimised as much as possible. All wild fish used in the experiments were released unharmed into the stream, and experimental females in the lab were returned to stock populations.

## References

Aronson, L. R. (1971). Further studies on orientation and jumping behavior in the gobiid fish, bathygobius soporator. Annals of the New York Academy of Sciences, 188(1), 378–392. https://doi.org/10.1111/J.1749-6632.1971.TB13110.X

Balda, R. P., & Kamil, A. C. (1992). Long-term spatial memory in clark’s nutcracker, Nucifraga columbiana. Animal Behaviour, 44(4), 761–769. https://doi.org/10.1016/S0003-3472(05)80302-1

Barki, A., Zion, B., Shapira, L., & Karplus, I. (2014). Using attraction to light to decrease cannibalism and increase fry production in guppy (Poecilia reticulata Peters) hatcheries. I: Phototactic reaction and light colour preference. Aquaculture Research, 45(8), 1295–1302. https://doi.org/10.1111/ARE.12070

Bates, D., Mächler, M., Bolker, B., & Walker, S. (2014). Fitting linear mixed-effects models using lme4. ArXiv Preprint 1406.5823.

Braithwaite, V. A., Armstrong, J. D., McAdam, H. M., & Huntingford, F. A. (1996). Can juvenile Atlantic salmon use multiple cue systems in spatial learning? Animal Behaviour, 51(6), 1409–1415. https://doi.org/10.1006/ANBE.1996.0144

Braithwaite, V. A., & Girvan, J. R. (2003). Use of water flow direction to provide spatial information in a small-scale orientation task. Journal of Fish Biology, 63(SUPPL. A), 74–83. https://doi.org/10.1111/J.1095-8649.2003.00218.X

Buo, C., Taylor, E., Dayal, P., Bartles, J., Christman, K., & Londraville, R. L. (2020a). Spatial mapping influences navigation in Entomacrodus striatus. Marine and Freshwater Behaviour and Physiology, 53(4), 193–201. https://doi.org/10.1080/10236244.2020.1785878

Buo, C., Taylor, E., Dayal, P., Bartles, J., Christman, K., & Londraville, R. L. (2020b). Spatial mapping influences navigation in Entomacrodus striatus. https://Doi.Org/10.1080/10236244.2020.1785878, 53(4), 193–201. https://doi.org/10.1080/10236244.2020.1785878

Burns, J. G., & Rodd, F. H. (2008a). Hastiness, brain size and predation regime affect the performance of wild guppies in a spatial memory task. Animal Behaviour, 76(3), 911–922. https://doi.org/10.1016/J.ANBEHAV.2008.02.017

Burns, J. G., & Rodd, F. H. (2008b). Hastiness, brain size and predation regime affect the performance of wild guppies in a spatial memory task. Animal Behaviour, 76, 911–922. https://doi.org/10.1016/j.anbehav.2008.02.017

Burt De Perera, T. (2004). Fish can encode order in their spatial map. Proceedings of the Royal Society of London. Series B: Biological Sciences, 271(1553), 2131–2134. https://doi.org/10.1098/RSPB.2004.2867

Chapman, B. B., Ward, A. J. W., & Krause, J. (2008). Schooling and learning: early social environment predicts social learning ability in the guppy, Poecilia reticulata. Animal Behaviour, 76(3), 923–929. https://doi.org/10.1016/J.ANBEHAV.2008.03.022

Cheng, K. (1986). A purely geometric module in the rat’s spatial representation. Cognition, 23(2), 149–178. https://doi.org/10.1016/0010-0277(86)90041-7

Croft, D. P., Albanese, B., Arrowsmith, B. J., Botham, M., Webster, M., & Krause, J. (2003). Sex-biased movement in the guppy (Poecilia reticulata). Oecologia, 137(1), 62–68. https://doi.org/10.1007/S00442-003-1268-6/FIGURES/2

Croft, D. P., Krause, J., & James, R. (2004). Social networks in the guppy (Poecilia reticulata). Proceedings of the Royal Society B: Biological Sciences, 271(SUPPL. 6), 516–519. https://doi.org/10.1098/rsbl.2004.0206

Darden, S. K., & Croft, D. P. (2008). Male harassment drives females to alter habitat use and leads to segregation of the sexes. Biology Letters, 4(5), 449–451. https://doi.org/10.1098/RSBL.2008.0308

Degen, J., Kirbach, A., Reiter, L., Lehmann, K., Norton, P., Storms, M., … Menzel, R. (2016). Honeybees Learn Landscape Features during Exploratory Orientation Flights. Current Biology, 26(20), 2800–2804. https://doi.org/10.1016/J.CUB.2016.08.013

Ehmer, B. (n.d.). Orientation in the Ant Paraponera clavata. Journal of Insect Behavior, 12(5). Retrieved from http://resolver.scholarsportal.info/resolve/08927553/v12i0005/711_oitapc.xml

Endler, J. A. (1992). Signals, signal conditions, and the direction of evolution. American Naturalist, 139(Suppl.). https://doi.org/10.1086/285308

Forward Jr, R. B., Horch, K. W., & Waterman, T. H. (1972). Visual orientation at the water surface by the teleost Zenarchopterus. The Biological Bulletin, 143(1), 112–126.

Fox, J., & Weisberg, S. (2018). An R companion to applied regression. Sage publications.

Goodyear, C. P., & Ferguson, D. E. (1969). Sun-compass orientation in the mosquitofish, Gambusia affinis. Animal Behaviour, 17(4), 636–640.

Gould, J. L. (1986). The Locale Map of Honey Bees: Do Insects Have Cognitive Maps? Science, 232(4752), 861–863. https://doi.org/10.1126/SCIENCE.232.4752.861

Hawkins, A. D., & Popper, A. N. (2018). Directional hearing and sound source localization by fishes. The Journal of the Acoustical Society of America, 144(6), 3329–3350.

Hölldobler, B. (1980). Canopy Orientation: A New Kind of Orientation in Ants. Science, 210(4465), 86–88. https://doi.org/10.1126/SCIENCE.210.4465.86

Hughes, R. N., & Blight, C. M. (2000). Two intertidal fish species use visual association learning to track the status of food patches in a radial maze. Animal Behaviour, 59(3), 613–621. https://doi.org/10.1006/ANBE.1999.1351

Kelly, D. M., Spetch, M. L., & Heth, C. D. (1998). Pigeons’ (Columba livia) Encoding of Geometric and Featural Properties of a Spatial Environment. Journal of Comparative Psychology, 112(3), 259–269. https://doi.org/10.1037/0735-7036.112.3.259

Kotrschal, A., Corral-Lopez, A., Amcoff, M., & Kolm, N. (2015). A larger brain confers a benefit in a spatial mate search learning task in male guppies. Behavioral Ecology, 26(2), 527–532. https://doi.org/10.1093/beheco/aru227

Landler, L., Ruxton, G. D., & Malkemper, E. P. (2019). The Hermans–Rasson test as a powerful alternative to the Rayleigh test for circular statistics in biology. BMC Ecology, 19(1), 1–8.

Lenth, R., Singmann, H., Love, J., Buerkner, P., & Herve, M. (2019). Package ‘emmeans.’

López, J. C., Broglio, C., Rodríguez, F., Thinus-Blanc, C., & Salas, C. (1999). Multiple spatial learning strategies in goldfish (Carassius auratus). Animal Cognition 1999 2:2, 2(2), 109–120. https://doi.org/10.1007/S100710050031

Lucon-Xiccato, T., & Bisazza, A. (2017a). Complex maze learning by fish. Animal Behaviour, 125, 69–75. https://doi.org/10.1016/J.ANBEHAV.2016.12.022

Lucon-Xiccato, T., & Bisazza, A. (2017b). Sex differences in spatial abilities and cognitive flexibility in the guppy. Animal Behaviour, 123, 53–60. https://doi.org/10.1016/j.anbehav.2016.10.026

Lucon-Xiccato, T., & Bisazza, A. (2017c). Sex differences in spatial abilities and cognitive flexibility in the guppy. Animal Behaviour, 123, 53–60. https://doi.org/10.1016/J.ANBEHAV.2016.10.026

Mazmanian, D. S., & Roberts, W. A. (1983). Spatial memory in rats under restricted viewing conditions. Learning and Motivation, 14(2), 123–139. https://doi.org/10.1016/0023-9690(83)90001-2

Miletto Petrazzini, M. E., Bisazza, A., Agrillo, C., & Lucon-Xiccato, T. (2017). Sex differences in discrimination reversal learning in the guppy. Animal Cognition, 20(6), 1081–1091. https://doi.org/10.1007/s10071-017-1124-4

Montgomery, J. C., Jeffs, A., Simpson, S. D., Meekan, M., & Tindle, C. (2006). Sound as an orientation cue for the pelagic larvae of reef fishes and decapod crustaceans. Advances in Marine Biology, 51, 143–196. https://doi.org/10.1016/S0065-2881(06)51003-X

Morris, R. G. M. (1981). Spatial localization does not require the presence of local cues. Learning and Motivation, 12(2), 239–260. https://doi.org/10.1016/0023-9690(81)90020-5

Noda, M., Gushima, K., & Kakuda, S. (1994). Local prey search based on spatial memory and expectation in the planktivorous reef fish, Chromis chrysurus (Pomacentridae). Animal Behaviour, 47(6), 1413–1422. https://doi.org/10.1006/ANBE.1994.1188

O’Keefe, J., & Conway, D. H. (1978). Hippocampal place units in the freely moving rat: why they fire where they fire. Experimental Brain Research, 31(4), 573–590. https://doi.org/10.1007/BF00239813

Odeh, M. (2002). Evaluation of the Effects of Turbulence on the Behavior of Migratory Fish, 2002 Final Report. US Geological Survey (USGS).

Odling-Smee, L., & Braithwaite, V. A. (2003). The role of learning in fish orientation. Fish and Fisheries, 4(3), 235–246.

Reese, E. S. (1989). Orientation behavior of butterflyfishes (family Chaetodontidae) on coral reefs: spatial learning of route specific landmarks and cognitive maps. Environmental Biology of Fishes 1989 25:1, 25(1), 79–86. https://doi.org/10.1007/BF00002202

Reznick, D., & Bryant, M. (2007). Comparative long-term mark-recapture studies of guppies (Poecilia reticulata): differences among high and low predation localities in growth and survival. Undefined.

Reznick, D. N., Butler IV, M. J., Rodd, F. H., & Ross, P. (1996). LIFE-HISTORY EVOLUTION IN GUPPIES (POECILIA RETICULATA) 6. DIFFERENTIAL MORTALITY AS A MECHANISM FOR NATURAL SELECTION. Evolution; International Journal of Organic Evolution, 50(4), 1651–1660. https://doi.org/10.1111/J.1558-5646.1996.TB03937.X

Rodriguez, F., Duran, E., Vargas, J. P., Torres, B., & Salas, C. (1994). Performance of goldfish trained in allocentric and egocentric maze procedures suggests the presence of a cognitive mapping system in fishes. Animal Learning & Behavior 1994 22:4, 22(4), 409–420. https://doi.org/10.3758/BF03209160

Salas, C., Broglio, C., Rodríguez, F., López, J. C., Portavella, M., & Torres, B. (1996). Telencephalic ablation in goldfish impairs performance in a ‘spatial constancy’ problem but not in a cued one. Behavioural Brain Research, 79(1–2), 193–200. https://doi.org/10.1016/0166-4328(96)00014-9

Seghers, B. H. (1973). Analysis of geographic variation in the antipredator adaptations of the guppy: Poecilia reticulata. University of British Columbia.

Simpson, S. D., Meekan, M., Montgomery, J., McCauley, R., & Jeffs, A. (2005). Homeward sound. Science, 308(5719), 221. https://doi.org/10.1126/SCIENCE.1107406

Soares, D., & Bierman, H. S. (2013). Aerial Jumping in the Trinidadian Guppy (Poecilia reticulata). PLOS ONE, 8(4), e61617. https://doi.org/10.1371/JOURNAL.PONE.0061617

Sovrano, V. A., Bisazza, A., & Vallortigara, G. (2002). Modularity and spatial reorientation in a simple mind: encoding of geometric and nongeometric properties of a spatial environment by fish. Cognition, 85(2), B51–B59. https://doi.org/10.1016/S0010-0277(02)00110-5

Suzuki, S., Augerinos, G., & Black, A. H. (1980). Stimulus control of spatial behavior on the eight-arm maze in rats. Learning and Motivation, 11(1), 1–18. https://doi.org/10.1016/0023-9690(80)90018-1

Teyke, T. (1989). Learning and remembering the environment in the blind cave fishAnoptichthys jordani. Journal of Comparative Physiology A 1989 164:5, 164(5), 655–662. https://doi.org/10.1007/BF00614508

Toates, F. M. (1980). Animal behaviour; a systems approach.

Tolimieri, N., Jeffs, A., & Montgomery, J. C. (2000). Ambient sound as a cue for navigation by the pelagic larvae of reel fishes. Marine Ecology Progress Series, 207, 219–224. https://doi.org/10.3354/MEPS207219

Tolman, E. C. (1948). Cognitive maps in rats and men. Psychological Review, 55(4), 189–208. https://doi.org/10.1037/H0061626

Warburton, K. (2003). Learning of foraging skills by fish. Fish and Fisheries, 4(3), 203–215. https://doi.org/10.1046/J.1467-2979.2003.00125.X

White, G. E., & Brown, C. (2014). A comparison of spatial learning and memory capabilities in intertidal gobies. Behavioral Ecology and Sociobiology, 68(9), 1393–1401. https://doi.org/10.1007/S00265-014-1747-2/FIGURES/5

