## Supplementary information for "Jumping out of trouble: Evidence for a cognitive map in guppies (*Poecilia reticulata*)"

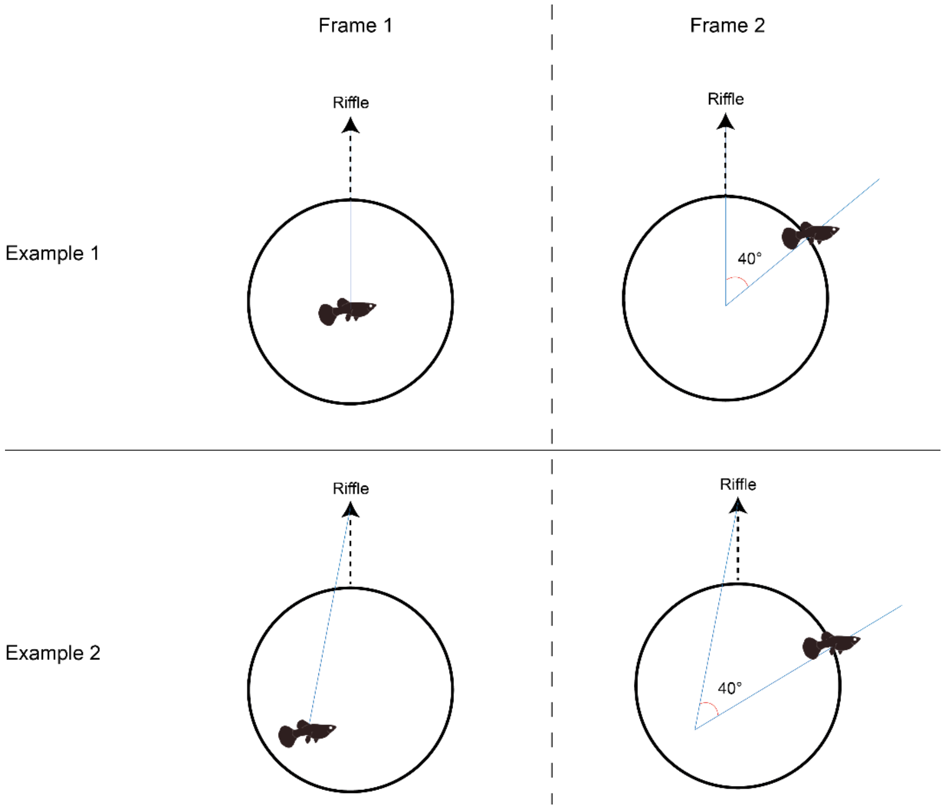


**Figure S1:** Schematic representations of the angle measurement with two different jump start positions used in Experiment 2 (Example 1 and Example 2). Frame 1 was selected when the fish started to protrude above the water surface. It was used to indicate the start position of the fish in reference to the riffle. Frame 2 frame was then selected when the fish crossed the edge of the cup. It was used to mark the position of the fish when crossing the edge of the cup. Subsequently, the angle was measured between the shortest path to the riffle centre and the centre point of the leaping fish when it crosses the edge of the cup.

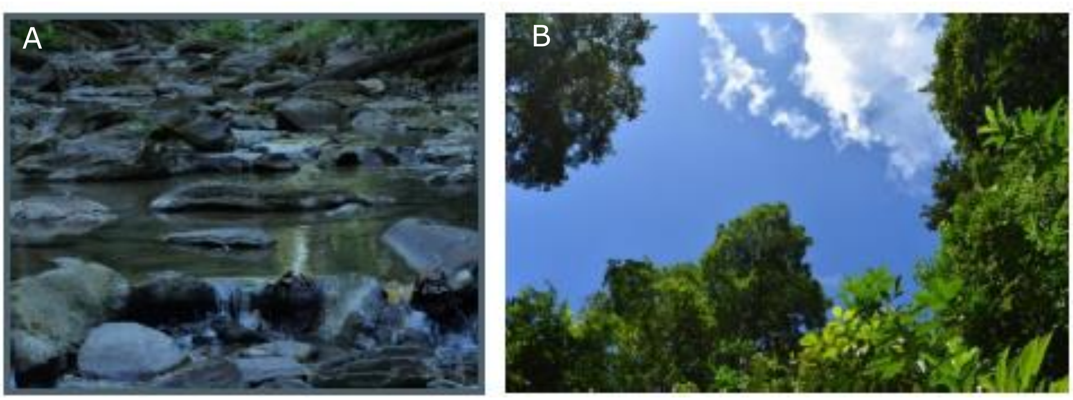


**Figure S2:** Photos of the test site in Experiment 2, including a riffle (A); and the canopy openness (B).
